# Determining the driving factors shaping genetic architecture of complex traits in recently admixed populations

**DOI:** 10.64898/2025.12.15.694350

**Authors:** Michelle Kim, Arun Durvasula, Xinjun Zhang

## Abstract

Understanding the genetic architecture of complex traits in admixed populations remains challenging due to heterogeneous genetic backgrounds and demographic histories. Mischaracterizing admixture can bias genetic association estimates and limit the generalizability of biomedical findings. Here, we systematically evaluate how evolutionary forces—including admixture, natural selection, and demographic history—jointly shape complex trait architecture and influence genome-wide association study (GWAS) outcomes using a simulation-based framework complemented by empirical analyses. We model five human admixture scenarios and vary the correlation between causal variant effect sizes and selection coefficients to reflect different trait–fitness relationships. This framework enables simulation of complex trait phenotypes with environmental variance, allowing comprehensive assessment of GWAS power and fine-mapping precision across evolutionary contexts. We find that GWAS power is strongly modulated by both genetic architecture and demographic history. Traits with weak coupling between fitness and effect size, such as anthropometric traits, exhibit higher GWAS power than traits under stronger negative selection, including early-onset diseases. Because rare variants contribute substantially to heritability yet are poorly captured by GWAS, bottlenecked populations with fewer rare variants show enhanced power. Despite large differences in GWAS power, fine-mapping precision remains relatively consistent across traits and populations, improving primarily in regions of high recombination. Empirical analyses of diverse cohorts, the All of Us Research Program, support these patterns. Our findings highlight how evolutionary and demographic forces shape the genetic basis of complex traits in admixed populations and underscore the need for tailored study designs to improve GWAS accuracy and fine-mapping performance in diverse cohorts.

## Introduction

Over the past two decades, genome-wide association studies (GWAS) have transformed the study of complex traits and diseases, yielding thousands of loci associated with phenotypes ranging from anthropometric measurements to disease susceptibility^1-4^. However, this progress has been accompanied by a persistent limitation, as the overwhelming majority of GWAS participants are of European ancestry^5-10^. Although initially driven by cohort availability, this imbalance now poses a major barrier to generalizing genetic findings across populations and to developing equitable applications in precision medicine^7,11^. As large-scale biobanks increasingly include individuals with more globally diverse backgrounds, including those with recent ancestry from multiple continental populations, there is a need to revisit assumptions underlying GWAS and to address the consequences of this long-standing representation gap.

Among globally diverse cohorts, admixed populations such as African American and Latino individuals present particularly informative but analytically complex settings. Their genomes are shaped by recent mixing between ancestrally distinct source populations, resulting in local ancestry mosaics and non-uniform linkage disequilibrium (LD) patterns^6,12^. Unlike in more homogeneous populations, the genomic architecture of admixed individuals reflects layered demographic histories including migration, bottlenecks, founder effects, as well as selective pressures that act differentially across ancestry tracts^13,14^. The resulting heterogeneity in allele frequencies and LD structure complicates association testing and challenges standard GWAS assumptions. In these contexts, even well-powered studies can yield biased or incomplete results unless population structure and local ancestry are properly accounted for.

Beyond demographic structure, genetic architecture itself – the distribution of causal variants, their allele frequencies, and effect sizes – is shaped by evolutionary forces. For traits under strong negative selection, such as early-onset genetic diseases, causal variants tend to be rare and exert relatively large effects^15-18^. In contrast, traits weakly coupled to fitness, like height or blood pressure, are typically influenced by numerous common variants of small effect^19^. Although these principles are well-established in evolutionary genetics, their impact on GWAS, particularly in admixed populations, remains incompletely understood. Prior studies have shown that GWAS power can vary with allele frequency and local ancestry, and that polygenic risk scores (PRS) derived in one ancestry often fail to transfer effectively across populations^6,20^. However, much of this work has examined one variable at a time. The joint influence of admixture, demography, selection, and trait architecture on association outcomes, especially in the context of rare variant detection and fine-mapping resolution, remains poorly characterized.

In this study, we systematically investigate how admixture events shape the genetic architecture of complex traits and influence GWAS performance, with the goal of establishing expectations and practical guidelines for future work. Specifically, we clarify when increasing cohort sizes in genetically diverse populations is likely or unlikely to improve power, and identify factors that most strongly affect the ability to identify true causal variants. We use forward-in-time simulations under a human demographic models, which allow us to track the true causal variants directly and evaluate which are successfully identified by GWAS and which are missed. These simulations model a range of evolutionary scenarios, including variation in recombination rate, selection intensity, admixture timing, and the degree of coupling between trait effects and fitness (Figure 1). By generating phenotypes under a spectrum of architectures we assess how these factors jointly shape GWAS power, heritability estimation, and fine-mapping resolution. To evaluate the relevance of these simulation-based patterns in real data, we complement our analysis with empirical GWAS results from the All of Us Research Program, a large-scale resource that includes participants from ancestrally diverse backgrounds. We compare simulated and empirical results across traits such as height and malignant neoplasms to evaluate whether evolutionary predictions are reflected in observed genetic patterns, including the greater contribution of common variants to polygenic traits and the enrichment of rare variants in disease-associated loci.

**Figure 1.**
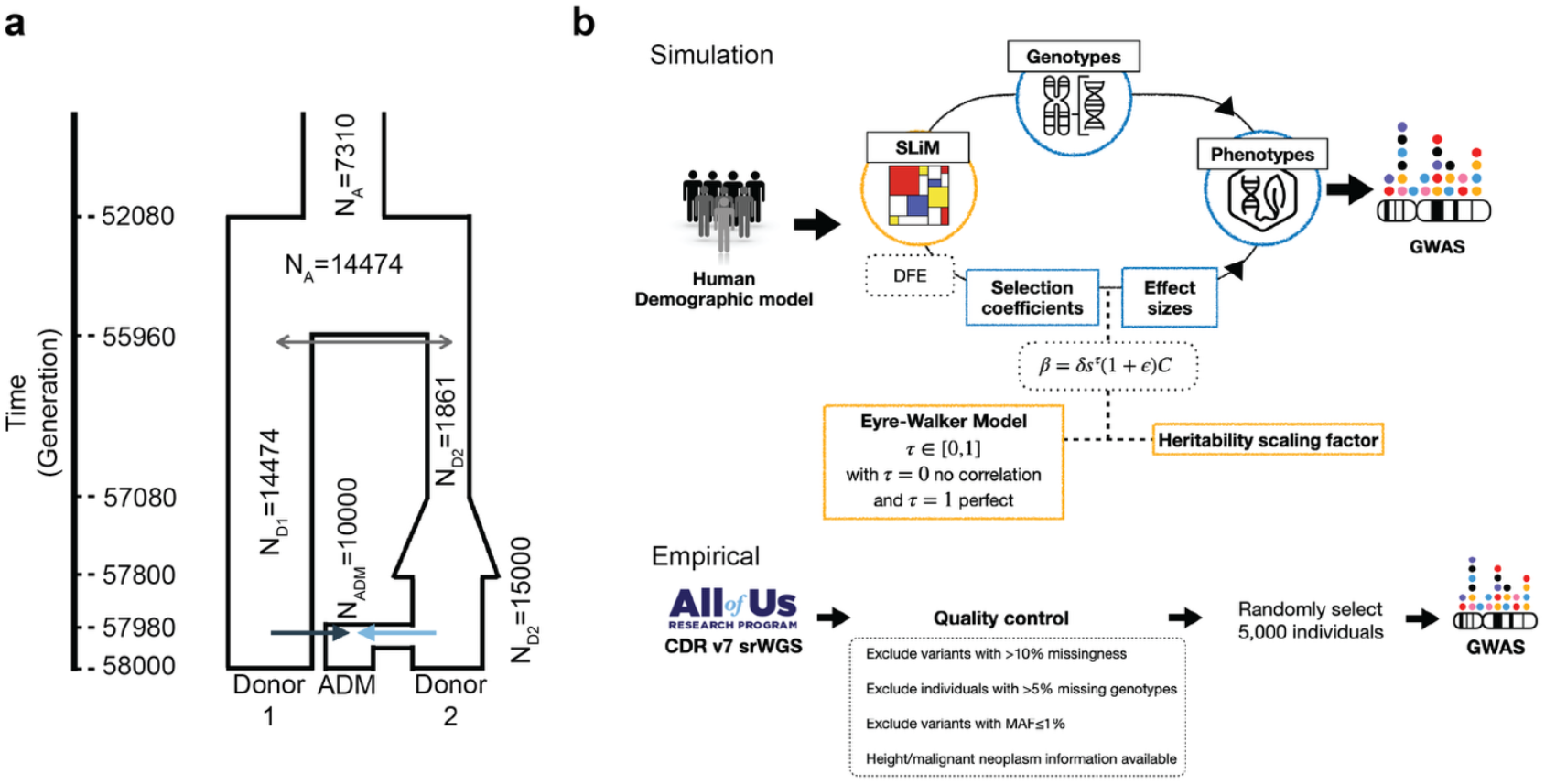
Workflow of the study. **a**. One of the five population demographic models explored. Parameters of the model as per Gravel et al. **b**. Workflow of the simulation and empirical data analysis.

## Materials and Methods

### Demographic models

To investigate how demographic history influences the genetic architecture of complex traits, we simulated a range of human demographic models based on the Gravel et al.^14^ Eurasian population history, incorporating migrations and admixture. In total, we designed five admixture models that reflect realistic admixture scenarios in empirical human populations, each with two sub-models with different majority of ancestry, yielding ten distinct simulation models (Supplementary Figure 1).

The five models vary mainly in terms of their admixture timing and demographic histories experienced by the ancestral populations (Donor 1 and Donor2). Model 1, which serves as the primary model for the analyses presented in the main text, represents a recent admixture scenario in which Donor 2 experiences a population bottleneck before the admixture event, following a prior phase of expansion. Model 2 captures an ancient admixture scenario that occurred >900 generations before sampling. In this model, Donor 2 undergoes an early bottleneck followed by continuous expansion until the present, where Donor 1 remains at a constant size. Model 3 mirrors Model 2 in its donor population histories but shifts the admixture event to a recent time point (20 generations before sampling), allowing us to isolate the effect of admixture timing on downstream genetic architecture. Model 4 builds on this framework by maintaining the same donor histories and admixture timing as Model 3 but introducing a smaller initial effective population size in the admixed group, modeling the impact of a founder effect and limited post-admixture expansion. Finally, Model 5 simulates a recent admixture event in which both donor populations experience simultaneous bottlenecks prior to admixture: Donor 1 undergoes a bottleneck for the first time, while Donor 2 experiences an additional bottleneck following its earlier expansion. For each of these five models, we implemented two sub-models to vary the admixture proportions: Model A and B. In model A, Donor 1 contributes 75% and Donor 2 contributes 25% to the admixed population (ADM-A). In model B, the proportions are reversed, with 25% from Donor 1 and 75% form Donor 2 (ADM-B). These models were designed to capture a range of realistic admixture histories, including differences in admixture timing, ancestry proportions, and population bottlenecks, providing a flexible framework for testing how demographic context shapes GWAS outcomes and trait architecture.

### Population demographic modeling and SLiM simulations

To model the genetic architecture of complex traits under a range of evolutionary and demographic scenarios, we used the forward-in-time simulation software SLiM^21^ v4.3. Simulations were grounded in one of the five admixture models incorporating population divergence, population size changes and migrations (Figure 1a). To examine the impact of demographic variables, we systematically varied parameters such as population size, migration rate, and recombination rate. Each simulation encompassed a 5 Mb genomic region with a mutation rate of 1.2x10^-8^ per base pair (bp) and a default recombination rate of 1x10^™8^ per bp, uniform across the simulated segment. When exploring the impact of recombination rate variation, we tested rates an order magnitude higher (1x10^-7^) or lower (1x10^-9^) than the default. Selection coefficients for de novo mutations were sampled from a gamma distribution with parameters estimated by Kim et al.^22^ (mean=-0.01026, *α*=0.186) allowing fitness effects to vary across the region. For each simulation replicate, we sampled 10,000 haploid genomes per population at the end of the run. if the population size was smaller than the target sample size, we increased it to the desired sample size after the scheduled endpoint and extended the simulation for 10 additional generations without introducing new mutations.

### Trait modeling and effect size assignment

As shown in Figure 1b, we modeled quantitative traits using an additive framework, drawing on the Eyre-Walker^16^ and extensions described by Lohmueller^17^. The effect size of each SNP (β) was defined as:

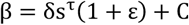

Here, δϵ{−1, 1} determines whether the variant increases or decreases the trait and is sampled with equal probability. The term ε∼N(0, 0.5) adds stochasticity to the effect size, and C is a scaling factor used to normalize trait heritability to 0.4. The selection coefficient (s) captures the fitness cost of the variant, τ ϵ [0, 1] modulates the correlation between effect size and fitness. We simulated three conditions: τ = 0 (no correlation), τ = 0.25 (weak correlation), and τ = 0.5 (moderate correlation), to reflect varying degrees of trait-fitness coupling.

Trait values for individuals were computed under an additive model:

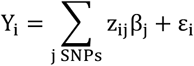

Where z_ij_ϵ{0,1,2} represents the genotype of individual at causal SNP *j*, β_j_ is the SNP’s effect size, and ε∼N(0, *V*_*E*_) is an environmental noise.

### GWAS and variant filtering

We conducted GWAS using linear regression on the simulated genotypes and phenotype values. To examine how allele frequency impacts GWAS performance, we performed two sets of analyses with different minor allele frequency (MAF) thresholds: (1) no threshold (all variants included) and (2) MAF≥5%. These thresholds allowed us to quantify how variant inclusion criteria affect downstream GWAS outcomes, particularly statistical power and fine-mapping precision. Power was defined as the proportion of true causal variants detected (sensitivity), and precision as the proportion of detected variants that were truly causal.

### Heritability and missing heritability estimation

To estimate the proportion of phenotypic variance attributable to genetic factors, we calculated narrow-sense heritability (*h*^*2*^) using the additive genetic variance framework given by

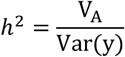

The additive genetic variance is 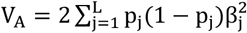 when there are *L* variants, where p_j_ is the allele frequency for variant *j* and β_j_ is the effect size of variant *j*.

### Polygenic score calculation

Polygenic scores were computed for each simulated individuals using effect size estimates obtained from the simulated GWAS. For each individual, we generated two types of polygenic scores: (1) matched PRS, in which GWAS summary statistics and the target individuals were drawn from the same population, and (2) transferred PRS, in which GWAS results from one population were applied to individuals from a different population. PRS were constructed using variants that surpassed genome-wide significance, defined by a Bonferroni-corrected p-value threshold, in the corresponding GWAS. For each individual *i*, the score was calculated as the sum of the genotype dosage weighted by the estimated effect size:

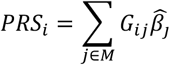

where *G*_*ij*_ ϵ {0, 1, 2} is the genotype dosage for variant *j* in individual 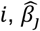 is the GWAS-estimated effect size of variant *j*, and *M* is the set of variants reaching genome-wide significance in the discovery GWAS.

### Empirical biobank analyses using All of Us

To evaluate whether patterns observed in simulated data extend to real-world populations, we analyzed genomic and phenotypic data from the All of Us Research Program (Controlled Tier Dataset v7)^23^. We restricted our analyses to participants with available short-read whole-genome sequencing (WGS) data and self-identified as African, White, or African American. Variant- and individual-level quality control was conducted using PLINK v2.0^24^. Variants with a missingness rate greater than 10% were excluded, as were individuals with more than 5% missing genotypes. We further applied MAF filter to ensure robustness in downstream association analyses.

Given the unknown trait-fitness relationship (τ) in real data, we selected two traits that likely differ in their selective profiles: standing height, assumed to have minimal coupling with fitness (τ ≈0), and malignant neoplasm, hypothesized to exhibit weak to moderate coupling ( τ > 0 ). To harmonize with the simulation study design, we randomly selected 5,000 individuals from each ancestry group to perform GWAS. For height, linear regression was used; for malignant neoplasm, a logistic regression framework was applied. Both models included sex and the first three principal components (PC1-PC3) as covariates to adjust for ancestry-related structure and potential confounding.

SNP-based heritability for both traits was estimated using the restricted maximum likelihood (GREML) approach as implemented in GCTA^25^. Individual-level phenotypes and covariates were used as inputs for variance component estimation.

PRS were computed using the pruning and thresholding (P+T) method, implemented via PLINK v2.0. LD pruning was performed using a *r*^*2*^ threshold of 0.1 in a sliding 250 kb window to ensure variant independence. A series of p-value thresholds were applied (5x10^-8^, 5x10^-7^, 5x10^-6^, 5x10^-4^, 5x10^-3^, 0.05, 0.1), and the threshold yielding the best predictive accuracy in a held-out subset was retained. Final PRS were calculated as the sum of risk allele dosages weighted by GWAS-derived effect sizes.

### Fine-mapping precision from UKB-PPP

To assess the fine-mapping precision of GWAS signals in empirical data, we leveraged protein quantitative trait loci (pQTL) results from the UK Biobank Pharma Proteomics Project (UKB-PPP), which includes high-resolution pQTL mapping across four biological panels: cardiometabolic, inflammation, neurology, and oncology^26^. Significant pQTLs variants were defined using the study-wide significance threshold of *p* < 1.7x10^-11^, accounting for multiple testing across both variants and proteins. These pQTL variants were merged with GWAS summary statistics from the UK Biobank Round 2, obtained from Neale Lab website (http://www.nealelab.is/uk-biobank/). For each phenotype, genome-wide significant loci were defined using a p-value threshold of *p*<5x10^-8^.

Fine-mapping precision was quantified as the proportion of GWAS-significant variants that overlapped with independently identified pQTLs from the corresponding biological panel:

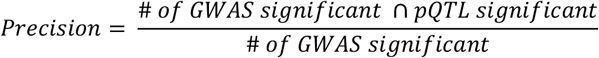

This calculation was performed separately for each panel, using curated phenotype-panel mappings based on biological relevance. The list of phenotypes assigned to each protein panel is provided in Supplementary Table1. Standard errors were obtained by jackknifing over loci, and these jackknife-derived estimates were used to construct error bars for the panel-specific precision values.

## Results

### Increasing the strength of selection reduces GWAS power while fine-mapping precision remains robust across populations and traits

Throughout this paper, we evaluate GWAS performance in terms of power, defined as the true positive rate (number of true causal among GWAS hits/total number of true causal), and fine mapping ability, defined as precision (number of true causal among GWAS hits/total number of GWAS hits). We assessed how GWAS power and fine-mapping precision are influenced by the coupling between selection and effect size, parameterized by *τ* (Figure 2). A low *τ* value (e.g. *τ*=0) indicates little correlation between effect size and the strength of negative selection, whereas a high *τ* value implies strong correlation between effect size and fitness. We further stratify and compare GWAS performance by the MAF thresholds and by the population in which the GWAS was performed. All simulation-based results and figures presented in the results section refer to Model 1 (recent admixture with Donor 2 bottlenecked), and results from the other four models are provided in the Supplementary Materials.

**Figure 2.**
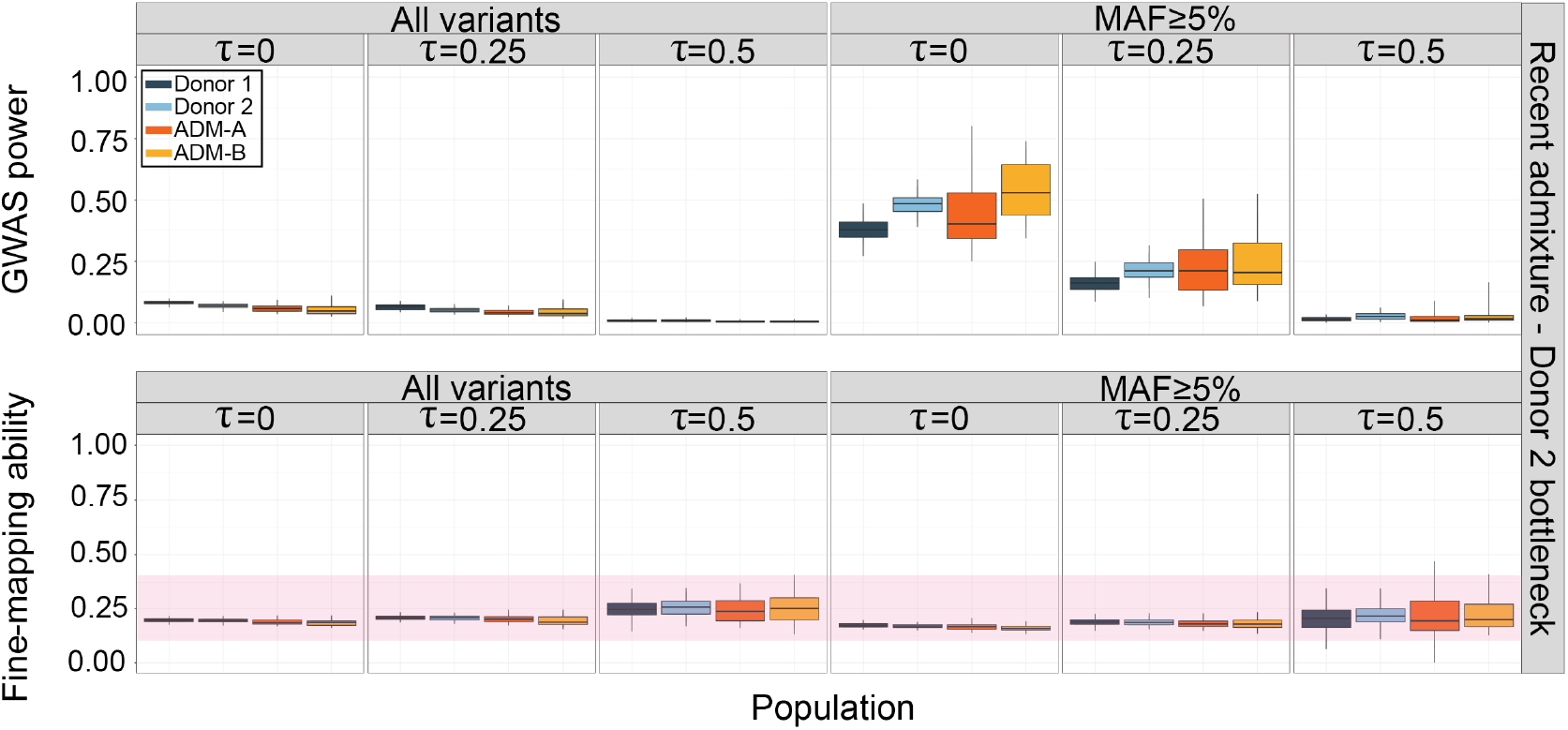
GWAS power and fine-mapping precision across populations and genetic architectures. GWAS power (top row. and fine-mapping precision (bottom row. across simulated populations indicated by color (dark blue: ancestral population 1; light blue: ancestral population 2; admixed population with more contribution from population 1: orange; admixed population with more contribution from population 2: yellow). Columns represent different values of the effect size-selection correlation parameter, with results shown for all variants (left. and for common variants (MAF≥5%; right). Each boxplot summarizes the distribution across replicate simulations per population.

First, we recapitulated the expected pattern that GWAS power depends strongly upon causal allele frequencies^27,28^. When all variants were included, GWAS power remained low across populations and *τ* values. In contrast, restricting the analysis to common variants (MAF≥5%) substantially increased power across all populations (Figure 2). This reflects the inherent difficulty of detecting rare variants: low allele frequencies reduce genetic variance, making it difficult for rare variants to achieve genome-wide significance.

Next, the extent to which selection shapes trait architecture also modulates GWAS power. When *τ* =0, indicating no correlation between effect size and selection, GWAS power was highest. In this regime, causal variants are distributed broadly across the allele frequency spectrum (Supplementary Figure 3). As *τ* increased to 0.5, power decreased across populations. When all variants were included in the GWAS, power declined by an average of about 6% across populations (ranging from 5% to 7%). When restricting the analysis to common variants, the decrease was more pronounced, averaging about 44% across populations (ranging from 37% to 50%). This reduction is consistent with the idea that large effect variants are efficiently removed by negative selection and only remain in low frequencies (Supplementary Figure 3), therefore are harder to detect using standard GWAS approaches under a strong correlation between effect size and selection. These patterns were consistent across populations, although the absolute power levels varied due to differences in allele frequency distributions and LD patterns.

When it comes to the fine-mapping precision, in contrast to GWAS power, fine-mapping precision remains surprisingly constant across all models we evaluated (Figure 2). Across all populations different *τ* values, with and without a MAF≥5% filter, we observe minimal change on GWAS precision as it remains relatively constant around 25%.

We note only a modest increase of precision when a trait is under stronger selection (high *τ* values). This slight increase likely reflects the fact that large-effect variants under strong negative selection are often rare and thus difficult for GWAS to detect, leaving fewer strong-effect variants for non-causal SNPs to tag. As a result, GWAS hits may stand out more clearly from the LD background, slightly improving fine mapping resolution. However, the magnitude of this effect was limited and is outweighed by the overall reduction of GWAS power for such traits.

### Ancestral demographic histories strongly shape GWAS power but not fine-mapping precision

Across all *τ* values, GWAS power in the admixed populations varied systematically with the demographic histories of the ancestral donor populations (Supplementary Figure 2). Relative to Model 1, Models 2 and 3, characterized by ancient or more recent admixture with continuous Donor 2 population growth, consistently showed significant lower power (Wilcoxon p<0.001 across *τ* values for both Pop3A and Pop3B). In contrast, Model 5, which incorporates bottlenecks in both Donor 1 and Donor 2, produced the highest power (p<10^-8^ across *τ* values), while Model 4, which adds a founder effect to an admixed population, did not differ significantly from Model 1 (p>0.3). This pattern consistent across both MAF strata and *τ* values, indicating that stronger drift and LD in bottlenecked donors enhance detection power in the admixed populations, whereas continuous growth in the donor populations reduces it.

In contrast, fine-mapping precision was largely stable across demographic models (Supplementary Figure 2). No significant differences were observed between Models 2–4 and Model 1 (adjusted p>0.3 across *τ* values and MAF strata), indicating that the accuracy of causal variant localization was robust to demographic variation in the ancestral donors. Only Model 5 exhibited marginally higher precision in a subset of cases (Pop 3A *τ*=0.25: p=0.002; Pop3A *τ*=0.5: p=0.0005; Pop3B *τ*=0.25 MAF≥5%: p=0.017), suggesting that the dual bottleneck scenario may slightly increase localization accuracy. Overall, while demographic history substantially influenced GWAS discovery power, its impact on fine-mapping precision was minimal.

### High recombination rate improves fine-mapping precision

We hypothesize that the relative stability of GWAS precision observed across models reflects the dominant role of LD block size in determining causal variant localization, with recombination acting as the primary force breaking correlations between causal and non-causal variants. To test this hypothesis, we examined the impact of recombination on GWAS performance by stratifying simulations into low (1x10^-9^), medium (1x10^-8^), and high (1x10^-7^) recombination rates, and jointly evaluating association power and fine-mapping precision.

Across all models, GWAS power declined with increasing recombination rate (Figure 3, solid line), a pattern most pronounced under moderate coupling between selection and effect size (*τ*=0.5). This reduction in power is consistent with the disruption of LD in high recombination regions, which lowers the likelihood that rare causal variants are tagged by nearby common variants and detected as GWAS hits. Conversely, extended LD in low recombination regions increases the likelihood that rare causal variants are indirectly captured, thereby enhancing detection power.

**Figure 3.**
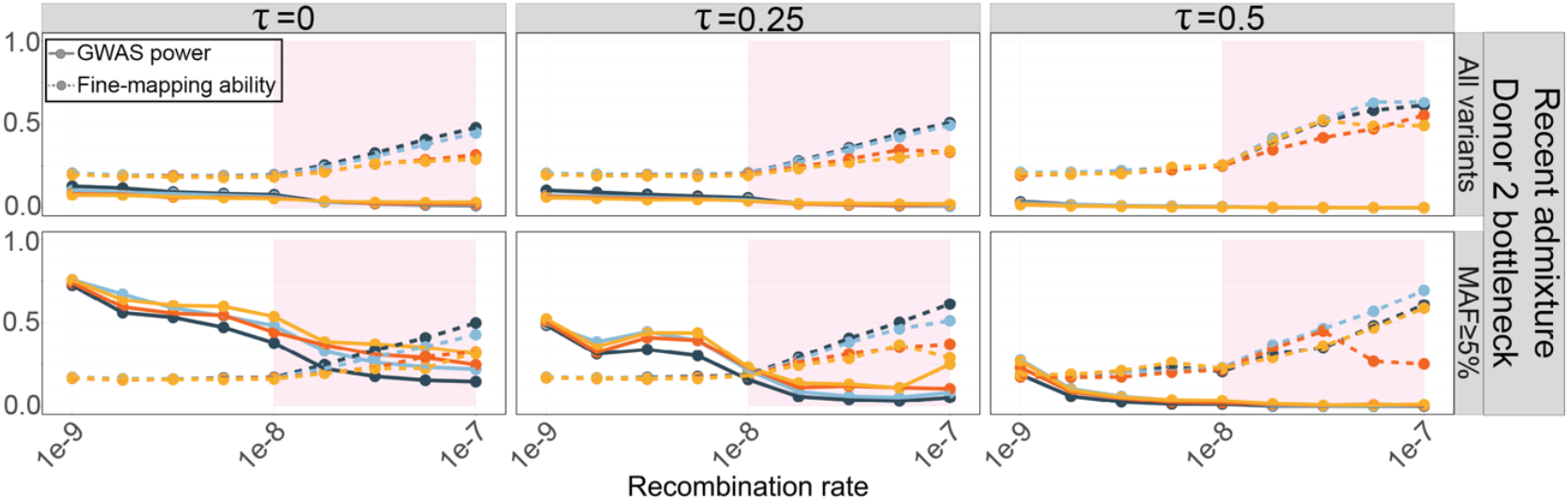
GWAS power and fine-mapping precision across recombination rates. Each line summarizes the mean GWAS power (solid. or fine-mapping precision (dashed. across replicated simulations for each population, shown across varying recombination rates. The pink shaded region in the background denotes the high recombination rate regime.

In contrast to association power, fine-mapping precision increased with recombination rate across all models (Figure 3, dashed line). This trend was robust across allele frequency thresholds and was most evident under *τ* =0.5, suggesting that more rapid LD decay in high-recombination regions improves localization of causal variants. As LD blocks shrink, fewer non-causal variants remain correlated with a causal variant, resulting in smaller credible sets and higher mapping resolution. This improvement was observed even when rare variants were included, indicating that elevated recombination systematically enhances fine-mapping accuracy rather than simply favoring common variants.

### Persistent missing heritability despite variation in trait architecture, recombination rates, and GWAS sample size

To evaluate how heritability estimation can vary across different traits and GWAS study designs, we quantified the proportion of total additive heritability explained by GWAS hits. Additive heritability is computed as 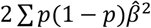, where *p* is the allele frequency and 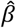 is the estimated effect size from GWAS.

Using simulations, we calculated the cumulative heritability recovered across allele frequency thresholds based on true effect sizes of causal variants and stratified results by population and selection effect size coupling parameter *τ* (Figure 4). Solid lines in Figure 4a indicate the total heritability attributable to all causal variants, while dashed lines represent the heritability explained by those causal variants detected through GWAS. This comparison isolates the role of statistical power in driving discrepancies between true and estimated heritability. To remove confounding introduced by recombination heterogeneity, analyses were performed at a fixed recombination rate of 1x10^-8^. When *τ* =0, representing traits whose effect sizes are independent of allele frequency, heritability was primarily driven by common variants. In contrast, under moderate coupling ( *τ* =0.5), causal variants were enriched for rarer alleles with larger effect sizes. Consequently, a substantial proportion of the heritability was attributable to low-frequency variants (e.g., MAF≤0.1%), and the gap between true and estimated heritability widened considerably. We further examined whether increasing GWAS sample size to 100,000 individuals could reduce missing heritability. Although greater sample size markedly improved heritability recovery, it did not fully close the gap between true and estimated heritability, with cumulative *h*^*2*^ from GWAS detected variants remaining below the ground truth (Supplementary Figure 5). Because GWAS power is limited for rare variants, most causal loci remained undetected, resulting in persistent missing heritability. These results indicate that incomplete heritability recovery reflects not only statistical power limitations but also fundamental features of underlying trait genetic architecture.

**Figure 4.**
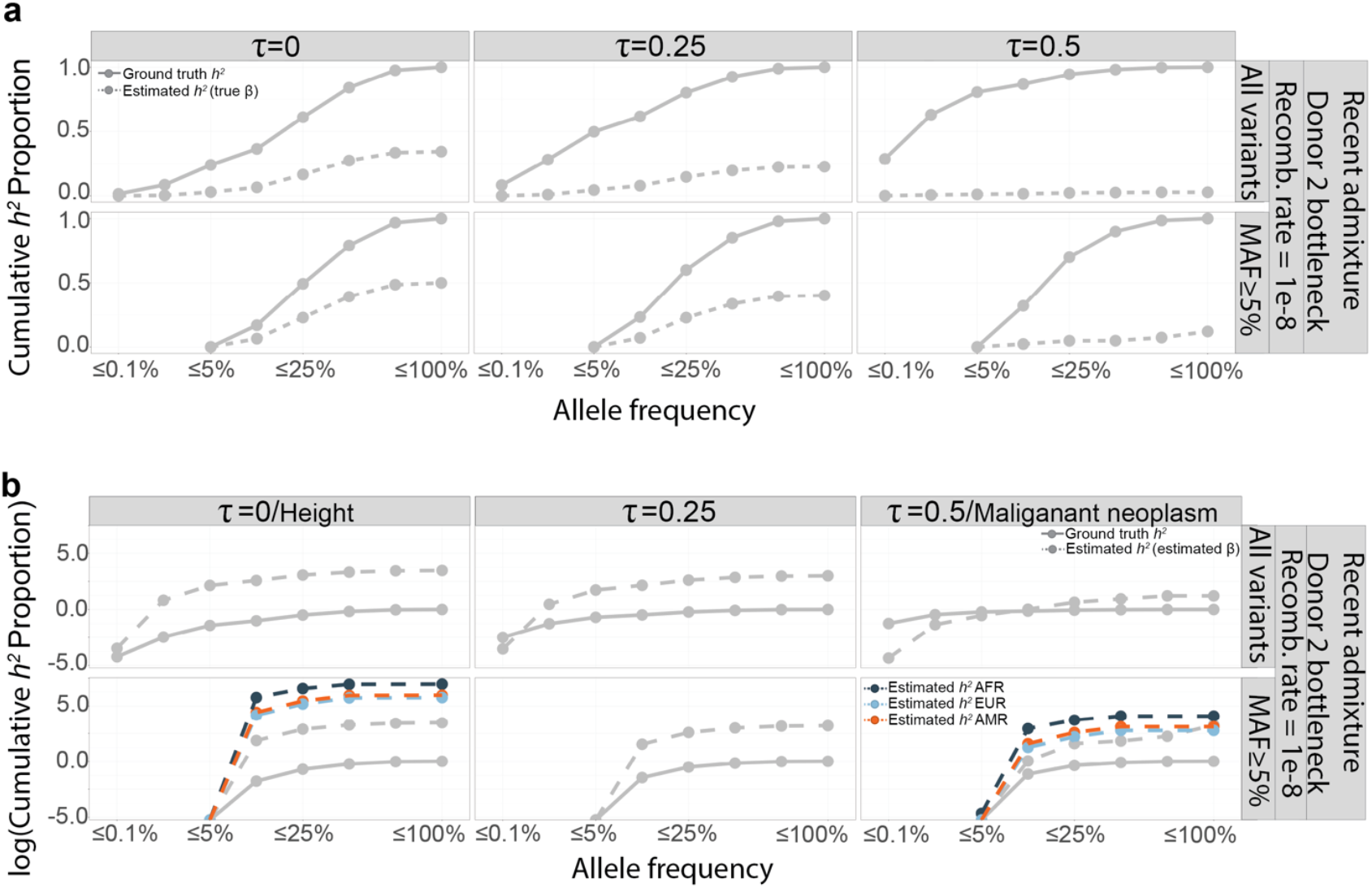
Cumulative contribution of variants to heritability across allele frequency bins. **a**. The solid lines represent the cumulative true heritability derived from all true causal variants (ground truth *h*^*2*^). The dashed lines represent the cumulative estimated heritability computed from genome-wide significant variants identified in GWAS and using true effect sizes (estimated *h*^*2*^–true β. **b**. Estimated heritability based on genome-wide significant variants with estimated effect sizes (grey dashed lines. compared against the ground truth heritability (solid grey lines). Colored lines are the estimated cumulative heritability proportion by each allele frequency bin from All of Us (Dark blue: AFR; Light blue: EUR; Orange: AMR).

Notably, when heritability was estimated using GWAS-significant variants with their estimated effect sizes, the cumulative *h*^*2*^ consistently exceeded the estimate obtained using the true effect sizes of the same variants (Figure 4b grey dashed line). This upward bias reflects the well-known winner’s curse: variants that achieve genome-wide significance tend to have effect sizes that are inflated by sampling error. Because heritability estimation depends on the effect size 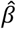, even modest upward bias in 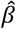 becomes magnified after squaring, leading to systematic overestimation of heritability.

### Population-matched GWAS maximizes polygenic risk prediction in homogeneous and admixed populations

We assessed the transferability of polygenic risk scores (PRS) by comparing predictive accuracy (using the coefficient of determination, *R*^*2*^) between simulated phenotypes and PRS constructed from GWAS conducted in different source populations (Figure 5). In all scenarios, PRS accuracy was highest when the GWAS and the target populations were matched, highlighting that population-matched GWAS consistently yields the most reliable predictions.

**Figure 5.**
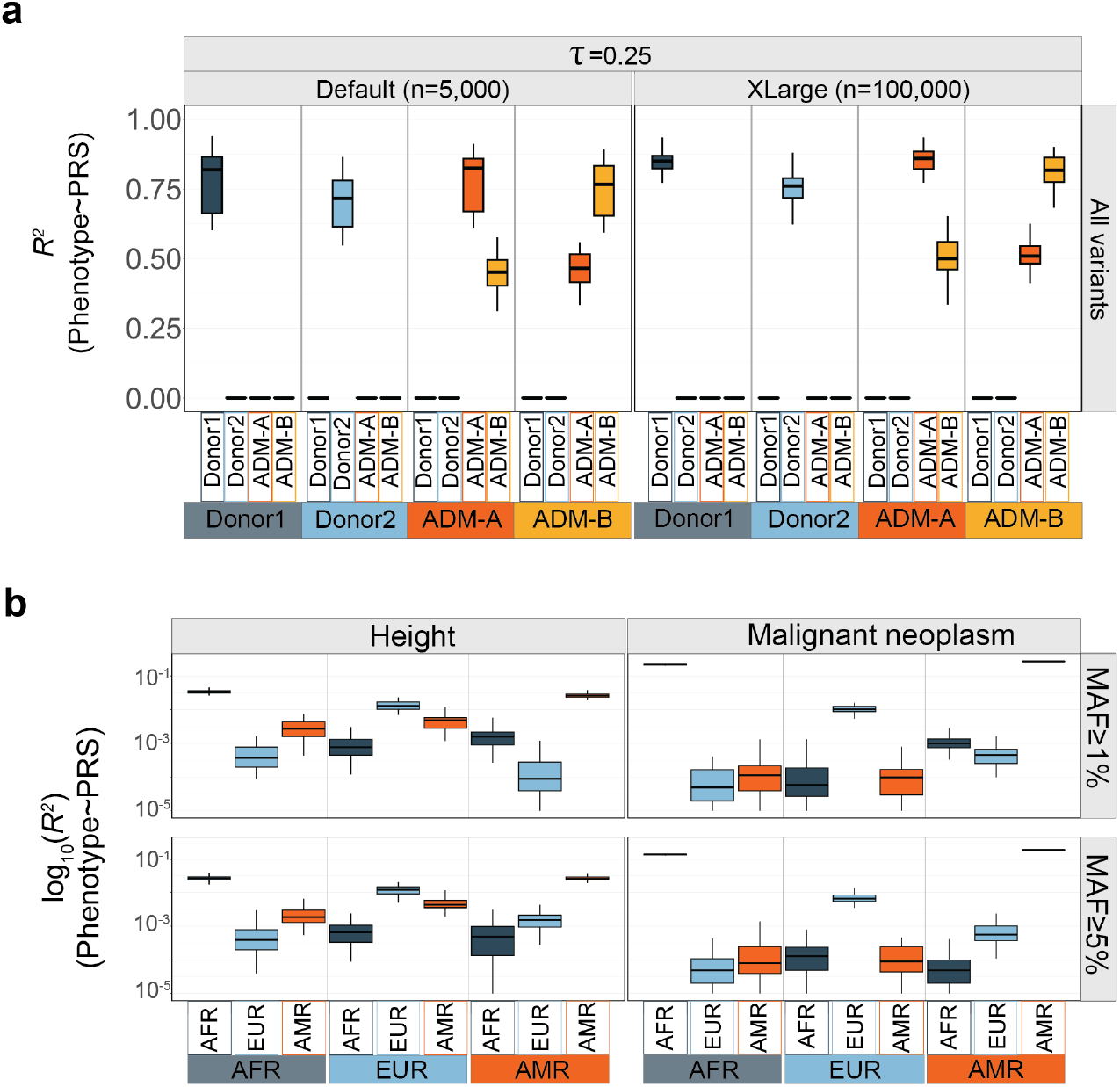
Phenotype ∼ PRS R^2^ across different GWAS and target population pairs under various trait-fitness coupling. **a**. Phenotypic variance explained (*R*^*2*^. by PRS across different GWAS and target population pairs under varying trait-fitness coupling (*τ*=0.25). Each boxplot represents the distribution of *R*^*2*^ values across simulation replicates, stratified by GWAS sample size (left: n=5,000; right: n=100,000). The x-axis denotes the target population in which PRS were applied, and boxplot colors indicate the GWAS population from which PRS were derived. Results are shown for all variants. **b**. Empirical PRS performance (log_10_(*R*^*2*^). across populations for two representative traits, height and malignant neoplasm, in the All of Us cohort. Each boxplot shows PRS accuracy when constructed from GWAS of the corresponding population (color-coded. and applied to target populations (x-axis). Results are stratified by MAF thresholds (top: MAF≥1%; bottom: MAF≥5%).

Under moderate coupling (*τ* =0.25), with a GWAS sample size of n=5,000 and no MAF filtering (Figure 5a), PRS applied within the same population achieved median *R*^*2*^ values between 0.75 and 0.90. In contrast, cross-population application of PRS produced substantially lower accuracy, with *R*^*2*^ values frequently below 0.4 and in some cases approaching zero when the ancestry divergence between GWAS and target populations was large. Increasing the GWAS sample size to n=100,000 improved accuracy overall but did not close the gap between matched and mismatched population pairs.

Importantly, this pattern also extended to admixed populations. Despite their heterogeneous ancestral backgrounds, PRS accuracy was consistently highest when GWAS was conducted directly in the target admixed population. Notably, transfers between differently admixed populations (e.g., ADM-A to ADM-B) achieved higher predictive accuracy than transfers from donor populations to admixed populations. This suggests that admixed-to-admixed PRS transfers preserves predictive signal more effectively than donor-to-admixed transfer, even when admixture proportions differ. Overall, our findings suggest that capturing the unique genetic architecture of the target population yields superior predictive performance than borrowing GWAS results from another population.

### Allele frequency filtering does not substantially impact fine-mapping precision in UK Biobank Pharma Proteomics Project (UKB-PPP)

In the previous section, we showed that fine-mapping precision, defined here as the precision of causal variants calls among GWAS hits, does not vary across different traits or GWAS study designs. Validating this in empirical data is challenging because the true causal variants underlying traits are generally unknown. To address this, we leveraged loci with functional evidence from protein quantitative trait loci (pQTLs) as a set of putative true positives to evaluate GWAS precision.

To assess whether allele frequency thresholds influence fine-mapping precision in empirical datasets, we analyzed GWAS summary statistics from UK Biobank alongside significant pQTLs identified in the UK Biobank Pharma Proteomics Project (UKB-PPP). Specifically, we calculated the proportion of GWAS-significant variants that were also independently identified as pQTLs, stratified by variant inclusion criteria (all variants vs. MAF≥5%) and by protein panel domain (cardiometabolic, inflammation, neurology, and oncology; Figure 6).

**Figure 6.**
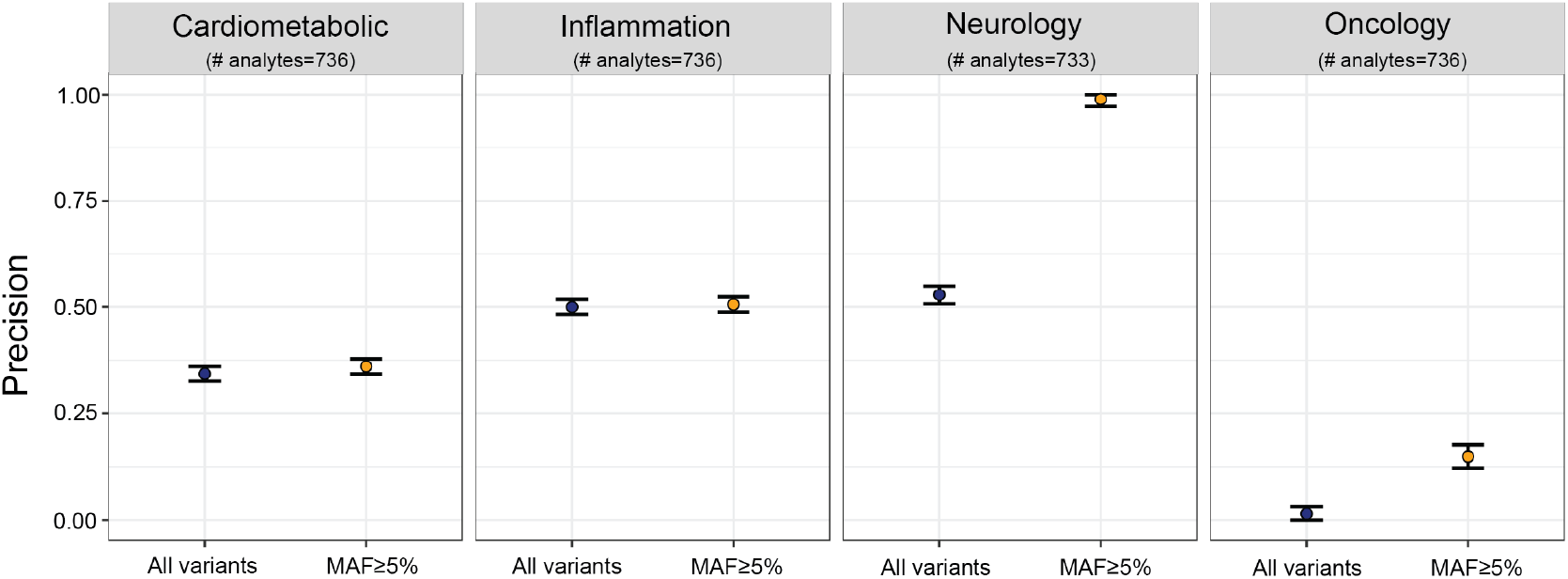
Fine-mapping precision across protein panels in empirical data. Precision calculated as the proportion of GWAS-significant variants that were also pQTL-significant (y-axis). Precision across four UKB-PPP protein panels under two variant inclusion criteria: all variants and MAF≥5% are shown.

Across domains, precision values varied, but applying a MAF filter had minimal effect on the overlap between GWAS and pQTL significant signals. For cardiometabolic and inflammation panels, precision remained stable regardless of MAF filtering, with only minimal increases of 0.02 (6%) and 0.01 (1%) respectively between unfiltered and MAF≥5% conditions. The neurology panel exhibited the highest fine-mapping precision overall, rising from 0.53 to 0.99 (an 87% increase) when restricting to common variants. In contrast, oncology showed the lowest precision under both conditions, though precision increased from 0.02 to 0.15 (a nearly tenfold improvement) when filtering for MAF≥5%. These results mirror our simulation-based findings, where filtering by allele frequency did not consistently improve fine-mapping resolution. Thus, while MAF thresholds may increase GWAS power, they do not substantially enhance the identification of true causal variants among statistically associated loci.

### All of Us data recapitulates upward bias in SNP-based heritability estimates

SNP-based heritability estimates from the All of Us populations showed cumulative patterns that closely mirrored those observed in the simulations (Figure 4b). Across allele frequency bins, the estimated curves rose more steeply than expected, reflecting consistent upward bias. This inflation reflects the tendency of variants that pass genome-wide significance thresholds to have overestimated effect sizes. As a result, cumulative heritability derived from GWAS-detected variants systematically exaggerates their contribution to trait phenotypic variance. These results suggest that part of the observed heritability in large-scale biobank studies reflects statistical estimation bias, consistent with the winner’s curse, rather than true underlying genetic signal.

### All of Us data confirms limited transferability of cross-population PRS

To empirically validate the simulation results that cross-population PRS performance is generally poor, we assessed PRS transferability in the All of Us cohort across three ancestry groups (AFR, EUR, and AMR) and two traits: height and malignant neoplasm (Figure 5b). For each ancestry group, we conducted GWAS and applied the resulting PRS to all three populations. Predictive accuracy was measured using the coefficient of determination *R*^*2*^ between PRS and phenotype, shown on the log_10_ scale. Each panel presents the PRS accuracy for a given GWAS-target pair, stratified by MAF threshold (MAF≥1% and MAF≥5%).

For height, predictive accuracy was consistently highest when GWAS and target populations were matched. PRS derived from European ancestry GWAS produced the highest *R*^*2*^ when applied to European individuals but showed marked reduction in performance when applied to AFR individuals, confirming limited transferability across divergent ancestries. This pattern was consistent across MAF thresholds, though restricting variants to MAF ≥ 5% led to a modest reduction in accuracy overall.

For malignant neoplasm, overall PRS predictive accuracy was lower across all populations, consistent with the trait’s more complex genetic architecture and lower heritability. Nonetheless, the same trend held: population-matched PRS yielded the highest predictive accuracy, while cross-population applications, particularly EUR-derived scores applied to AFR individuals, showed the steepest performance declines. These empirical findings mirror our simulation results and reinforce the conclusion that PRS transferability across populations is limited, underscoring the importance of performing GWAS directly in the target population, including admixed populations.

## Discussion

Understanding the factors that shape GWAS performance across diverse populations is essential for improving risk prediction in globally representative cohorts. In this study, we provide a comprehensive simulation-based framework to address the long-standing question of how evolutionary processes, including admixture, shape the genetic architecture and complex traits and influence the performance of association mapping methods such as GWAS. By explicitly tracking true causal variants in simulations, we were able to distinguish variants detected by GWAS from those that remain undetected, enabling a systematic evaluation of the factors governing the distribution of causal variants in populations. Through this approach, we examined how demographic history (including population size changes and admixture events), genomic properties such as recombination rate, and trait-specific genetic architecture of traits characterized by the relationship between selection and effect size, jointly influence GWAS power, fine-mapping precision, missing heritability, and polygenic risk score portability in admixed populations. These simulation-based predictions were further supported by empirical analyses of data from the UK Biobank and the All of Us Research Program, allowing us to directly compare theoretical expectations with real-world genetic datasets.

Our results show that GWAS power is strongly shaped by the interplay between trait architecture and local genomic features. When causal effect sizes were independent of allele frequency (i.e., *τ*=0), power was highest and heritability was primarily attributable to common variants. In contrast, under modest coupling between selection and effect size (*τ*=0.25-0.5), power declined sharply, particularly when rare, strongly selected variants contributed to the trait. This reduction in power directly led to the underestimation of heritability: as rare variants explained a larger share of genetic variance, the gap between true and GWAS-estimated heritability widened, reflecting a form of missing heritability that GWAS is intrinsically unable to recover due to its extreme underpower in calling low-frequency causal loci. These findings highlight the need for developing novel approach beyond standard GWAS to capture the contribution of rare variants.

Importantly, while MAF-based variant filtering improved GWAS power, it did not meaningfully enhance fine-mapping precision, consistent with prior work suggesting that LD structure and tagging efficiency, rather than allele frequency alone, govern resolution. Notably, even if rare variants are difficult to detect, they are not impossible to be called: we show that under a uniform recombination rate of 1x10^-8^, GWAS precision remained stable at 25% across both common and rare variants. As recombination rate increased, GWAS power declined rapidly due to faster LD decay and reduced tagging. In contrast, fine mapping precision improved as LD blocks become smaller, enhancing localization accuracy. This tradeoff between declining power and increasing precision suggests a potential pathway for identifying true causal variants, including rare ones, even if when discovery is limited to a modest number of loci. Such high-confidence variant sets may provide a foundation for developing new methods for rare variant discovery, which remains a long-standing challenge in complex trait genetics. We further show that this tradeoff was most pronounced under architectures shaped by selection, where causal variants tend to be rare and poorly tagged. These findings underscore the importance of considering recombination context when interpreting association signals and prioritizing causal variants, particularly for traits influenced by selection.

Analyses of extra-large GWAS (N=100,000) showed that increasing sample size only incrementally improved power and did not enhance fine-mapping precision. Larger cohorts improved detection of common variants but provided limited gains for localizing causal variants, as precision remained constrained by LD structure. PRS performance showed a similar pattern: accuracy increased slightly with larger discovery GWAS but remained highly dependent on matching the discovery and target population. Predictive accuracy was highest within the same population and lowest in African ancestry groups, reflecting reduced tagging and higher allele frequency differentiation. These results indicate that scaling GWAS alone cannot resolve the limitations of power, precision, and portability across populations.

Patterns observed in simulations were recapitulated in empirical data. Cross-population PRS analyses in the All of Us cohort revealed strong ancestry dependence, with predictive accuracy highest when the GWAS and target populations matched. PRS portability was lowest for AFR individuals, consistent with reduced tagging and higher allele frequency differentiation. For traits such as malignant neoplasm, which are likely influenced by rare or ancestry-specific variants, PRS accuracy was uniformly low across populations. In fine-mapping analyses based on pQTL and GWAS concordance in UK Biobank, precision varied by trait domain but remained largely unchanged with MAF filtering, mirroring the simulation-based observation that filtering alone does not improve localization. Among the protein panels analyzed, the oncology domain exhibited notably lower fine-mapping precision compared to cardiometabolic, inflammation, and neurology panels. This may reflect biological and methodological differences across protein traits. Many oncology-associated proteins are upregulated in disease states but may be minimally expressed or undetectable in healthy individuals, reducing statistical power for pQTL detection in population-scale biobanks^29^. Additionally, cancer-related proteins may be influenced by somatic mutations or context-specific regulatory mechanisms not captured in germline variation-based GWAS, further limiting overlap with pQTL signals^30,31^. These factors likely contribute to the reduced concordance between GWAS and pQTL signals in the oncology panel and underscore the importance of trait-specific considerations when evaluating fine-mapping performance using orthogonal datasets.

Together, these findings highlight key limitations of standard GWAS designs in recovering the genetic architecture of complex traits, especially in the presence of rare variants and population structure. Our results emphasize that both recombination landscape and evolutionary coupling between fitness and trait effect shape GWAS outcomes, and that these forces must be accounted for to improve power, portability, and resolution. As large-scale biobanks continue to expand in diversity and scale, incorporating selection-aware, LD-aware, and rare variant-sensitive models will be critical for equitable and accurate genetic discovery.

## Acknowledgement

We gratefully acknowledge *All of Us* participants for their contributions, without whom this research would not have been possible. We also thank the National Institutes of Health’s [<u>*All of Us* Research Program</u>](https://allofus.nih.gov/) for making available the participant data [and/or samples and/or cohort] examined in this study.

## References

1. Bycroft, C., Freeman, C., Petkova, D., Band, G., Elliott, L.T., Sharp, K., Motyer, A., Vukcevic, D., Delaneau, O., O’Connell, J., et al. (2018). The UK Biobank resource with deep phenotyping and genomic data. Nature 562, 203–209. 10.1038/s41586-018-0579-z.

2. Consortium, U.K., Walter, K., Min, J.L., Huang, J., Crooks, L., Memari, Y., McCarthy, S., Perry, J.R., Xu, C., Futema, M., et al. (2015). The UK10K project identifies rare variants in health and disease. Nature 526, 82–90. 10.1038/nature14962.

3. Visscher, P.M., Wray, N.R., Zhang, Q., Sklar, P., McCarthy, M.I., Brown, M.A., and Yang, J. (2017). 10 Years of GWAS Discovery: Biology, Function, and Translation. Am J Hum Genet 101, 5–22. 10.1016/j.ajhg.2017.06.005.

4. Wellcome Trust Case Control, C. (2007). Genome-wide association study of 14,000 cases of seven common diseases and 3,000 shared controls. Nature 447, 661–678. 10.1038/nature05911.

5. Kim, M.S., Patel, K.P., Teng, A.K., Berens, A.J., and Lachance, J. (2018). Genetic disease risks can be misestimated across global populations. Genome Biol 19, 179. 10.1186/s13059-018-1561-7.

6. Martin, A.R., Gignoux, C.R., Walters, R.K., Wojcik, G.L., Neale, B.M., Gravel, S., Daly, M.J., Bustamante, C.D., and Kenny, E.E. (2017). Human Demographic History Impacts Genetic Risk Prediction across Diverse Populations. Am J Hum Genet 100, 635–649. 10.1016/j.ajhg.2017.03.004.

7. Martin, A.R., Kanai, M., Kamatani, Y., Okada, Y., Neale, B.M., and Daly, M.J. (2019). Clinical use of current polygenic risk scores may exacerbate health disparities. Nat Genet 51, 584–591. 10.1038/s41588-019-0379-x.

8. Mostafavi, H., Harpak, A., Agarwal, I., Conley, D., Pritchard, J.K., and Przeworski, M. (2020). Variable prediction accuracy of polygenic scores within an ancestry group. Elife 9. 10.7554/eLife.48376.

9. Ragsdale, A.P., Nelson, D., Gravel, S., and Kelleher, J. (2020). Lessons Learned from Bugs in Models of Human History. Am J Hum Genet 107, 583–588. 10.1016/j.ajhg.2020.08.017.

10. Scutari, M., Mackay, I., and Balding, D. (2016). Using Genetic Distance to Infer the Accuracy of Genomic Prediction. PLoS Genet 12, e1006288. 10.1371/journal.pgen.1006288.

11. Popejoy, A.B., and Fullerton, S.M. (2016). Genomics is failing on diversity. Nature 538, 161–164. 10.1038/538161a.

12. Wojcik, G.L., Graff, M., Nishimura, K.K., Tao, R., Haessler, J., Gignoux, C.R., Highland, H.M., Patel, Y.M., Sorokin, E.P., Avery, C.L., et al. (2019). Genetic analyses of diverse populations improves discovery for complex traits. Nature 570, 514–518. 10.1038/s41586-019-1310-4.

13. Bryc, K., Durand, E.Y., Macpherson, J.M., Reich, D., and Mountain, J.L. (2015). The genetic ancestry of African Americans, Latinos, and European Americans across the United States. Am J Hum Genet 96, 37–53. 10.1016/j.ajhg.2014.11.010.

14. Gravel, S., Henn, B.M., Gutenkunst, R.N., Indap, A.R., Marth, G.T., Clark, A.G., Yu, F., Gibbs, R.A., Genomes, P., and Bustamante, C.D. (2011). Demographic history and rare allele sharing among human populations. Proc Natl Acad Sci U S A 108, 11983–11988. 10.1073/pnas.1019276108.

15. Durvasula, A., and Lohmueller, K.E. (2021). Negative selection on complex traits limits phenotype prediction accuracy between populations. Am J Hum Genet 108, 620–631. 10.1016/j.ajhg.2021.02.013.

16. Eyre-Walker, A. (2010). Evolution in health and medicine Sackler colloquium: Genetic architecture of a complex trait and its implications for fitness and genome-wide association studies. Proc Natl Acad Sci U S A 107 Suppl 1, 1752–1756. 10.1073/pnas.0906182107.

17. Lohmueller, K.E. (2014). The impact of population demography and selection on the genetic architecture of complex traits. PLoS Genet 10, e1004379. 10.1371/journal.pgen.1004379.

18. Zeng, J., de Vlaming, R., Wu, Y., Robinson, M.R., Lloyd-Jones, L.R., Yengo, L., Yap, C.X., Xue, A., Sidorenko, J., McRae, A.F., et al. (2018). Signatures of negative selection in the genetic architecture of human complex traits. Nat Genet 50, 746–753. 10.1038/s41588-018-0101-4.

19. Boyle, E.A., Li, Y.I., and Pritchard, J.K. (2017). An Expanded View of Complex Traits: From Polygenic to Omnigenic. Cell 169, 1177–1186. 10.1016/j.cell.2017.05.038.

20. Atkinson, E.G., Maihofer, A.X., Kanai, M., Martin, A.R., Karczewski, K.J., Santoro, M.L., Ulirsch, J.C., Kamatani, Y., Okada, Y., Finucane, H.K., et al. (2021). Tractor uses local ancestry to enable the inclusion of admixed individuals in GWAS and to boost power. Nat Genet 53, 195–204. 10.1038/s41588-020-00766-y.

21. Haller, B.C., and Messer, P.W. (2023). SLiM 4: Multispecies Eco-Evolutionary Modeling. Am Nat 201, E127–E139. 10.1086/723601.

22. Kim, B.Y., Huber, C.D., and Lohmueller, K.E. (2017). Inference of the Distribution of Selection Coefficients for New Nonsynonymous Mutations Using Large Samples. Genetics 206, 345–361. 10.1534/genetics.116.197145.

23. All of Us Research Program, I., Denny, J.C., Rutter, J.L., Goldstein, D.B., Philippakis, A., Smoller, J.W., Jenkins, G., and Dishman, E. (2019). The “All of Us” Research Program. N Engl J Med 381, 668–676. 10.1056/NEJMsr1809937.

24. Chang, C.C., Chow, C.C., Tellier, L.C., Vattikuti, S., Purcell, S.M., and Lee, J.J. (2015). Second-generation PLINK: rising to the challenge of larger and richer datasets. Gigascience 4, 7. 10.1186/s13742-015-0047-8.

25. Yang, J., Lee, S.H., Goddard, M.E., and Visscher, P.M. (2011). GCTA: a tool for genomewide complex trait analysis. Am J Hum Genet 88, 76–82. 10.1016/j.ajhg.2010.11.011.

26. Sun, B.B., Chiou, J., Traylor, M., Benner, C., Hsu, Y.H., Richardson, T.G., Surendran, P., Mahajan, A., Robins, C., Vasquez-Grinnell, S.G., et al. (2023). Plasma proteomic associations with genetics and health in the UK Biobank. Nature 622, 329–338. 10.1038/s41586-023-06592-6.

27. Park, J.H., Gail, M.H., Weinberg, C.R., Carroll, R.J., Chung, C.C., Wang, Z., Chanock, S.J., Fraumeni, J.F., Jr., and Chatterjee, N. (2011). Distribution of allele frequencies and effect sizes and their interrelationships for common genetic susceptibility variants. Proc Natl Acad Sci U S A 108, 18026–18031. 10.1073/pnas.1114759108.

28. Lee, S., Abecasis, G.R., Boehnke, M., and Lin, X. (2014). Rare-variant association analysis: study designs and statistical tests. Am J Hum Genet 95, 5–23. 10.1016/j.ajhg.2014.06.009.

29. Zhang, J., Dutta, D., Kottgen, A., Tin, A., Schlosser, P., Grams, M.E., Harvey, B., Consortium, C.K., Yu, B., Boerwinkle, E., et al. (2022). Plasma proteome analyses in individuals of European and African ancestry identify cis-pQTLs and models for proteomewide association studies. Nat Genet 54, 593–602. 10.1038/s41588-022-01051-w.

30. Papier, K., Atkins, J.R., Tong, T.Y.N., Gaitskell, K., Desai, T., Ogamba, C.F., Parsaeian, M., Reeves, G.K., Mills, I.G., Key, T.J., et al. (2024). Identifying proteomic risk factors for cancer using prospective and exome analyses of 1463 circulating proteins and risk of 19 cancers in the UK Biobank. Nat Commun 15, 4010. 10.1038/s41467-024-48017-6.

31. Sun, B.B., Maranville, J.C., Peters, J.E., Stacey, D., Staley, J.R., Blackshaw, J., Burgess, S., Jiang, T., Paige, E., Surendran, P., et al. (2018). Genomic atlas of the human plasma proteome. Nature 558, 73–79. 10.1038/s41586-018-0175-2.

