## Supplementary files for "Determining the driving factors shaping genetic architecture of complex traits in recently admixed populations"

### Supplementary Figures

**Figure S1)** Demographic models used in simulations with population size parameters and generation times indicated. Across the demographic scenarios, admixture timing, population size, and bottleneck history are explored.

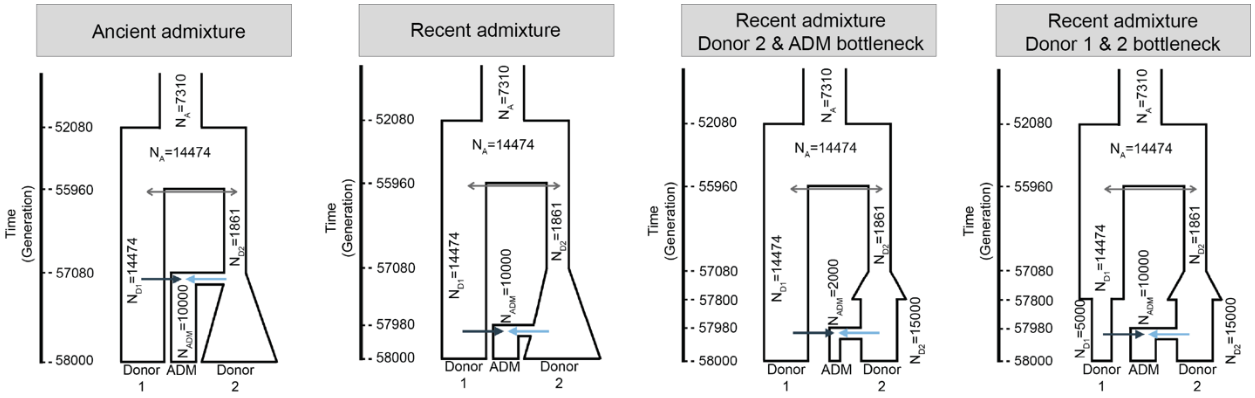

**Figure S2)** GWAS power and fine-mapping ability across populations and genetic architectures.

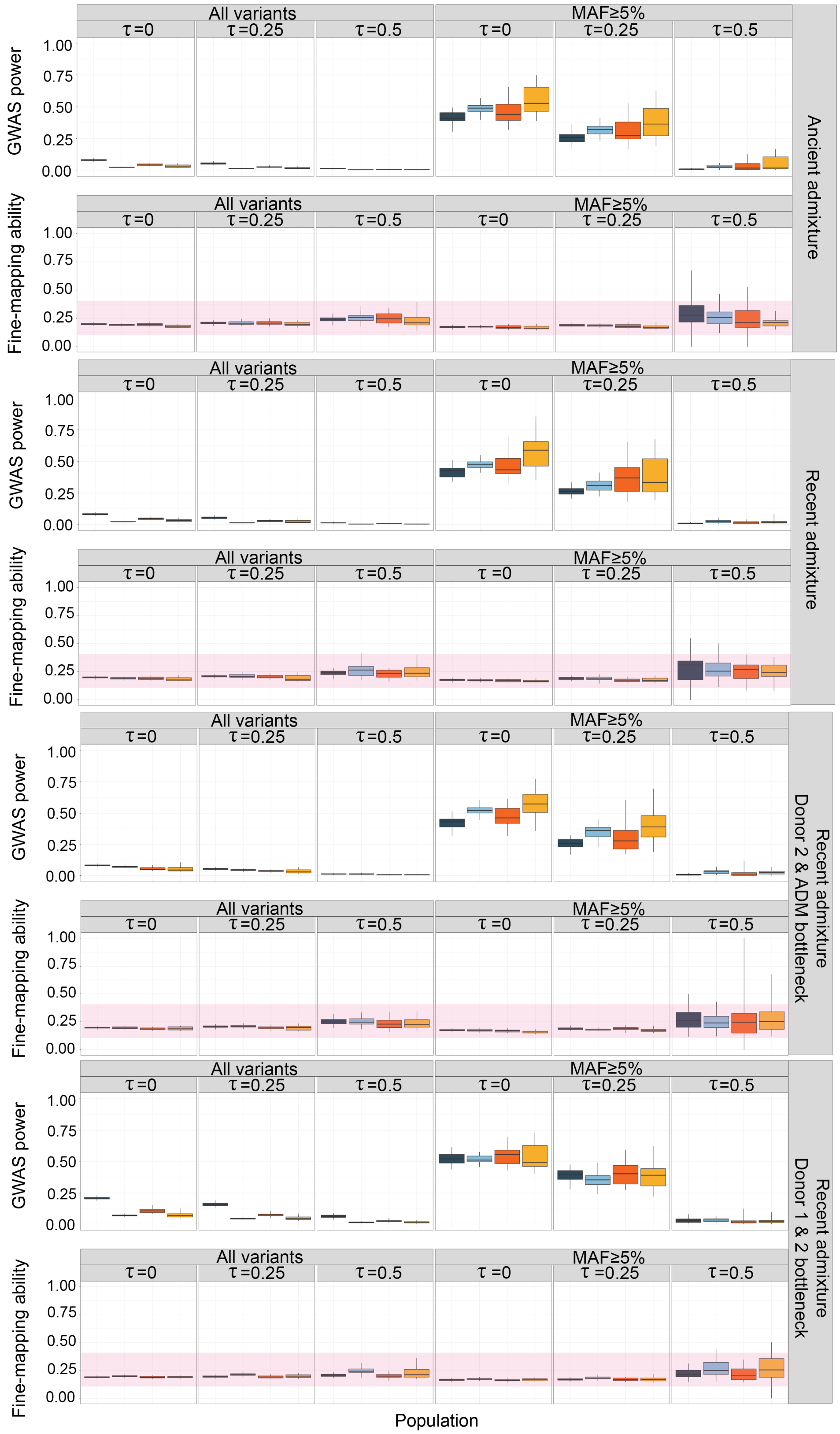

**Figure S3)** Relationship between MAF and effect size across populations in simulated and empirical data.
Top row: Simulated data showing effect sizes of causal variants under two trait architectures: $\tau$=0 (left) and $\tau$=0.5 (right). Columns correspond to different populations: Donor 1, Donor 2, ADM-A, and ADM-B.
Bottom row: Empirical GWAS summary statistics from the All of Us Research Program for two traits – height (left) and malignant neoplasm (right) – in AFR, EUR, and AMR populations.

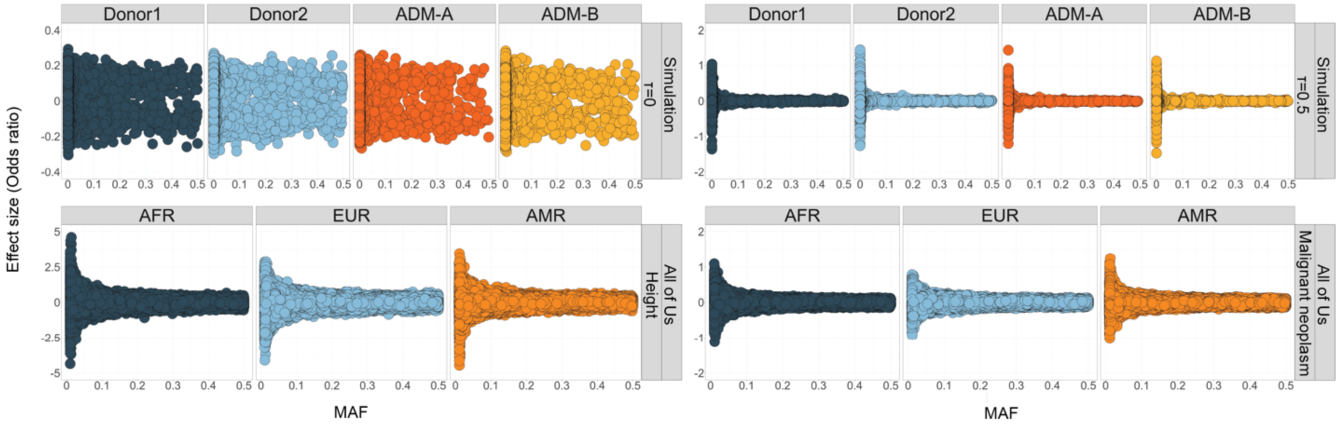

**Figure S4) GWAS power and fine-mapping ability (precision) by recombination rates**

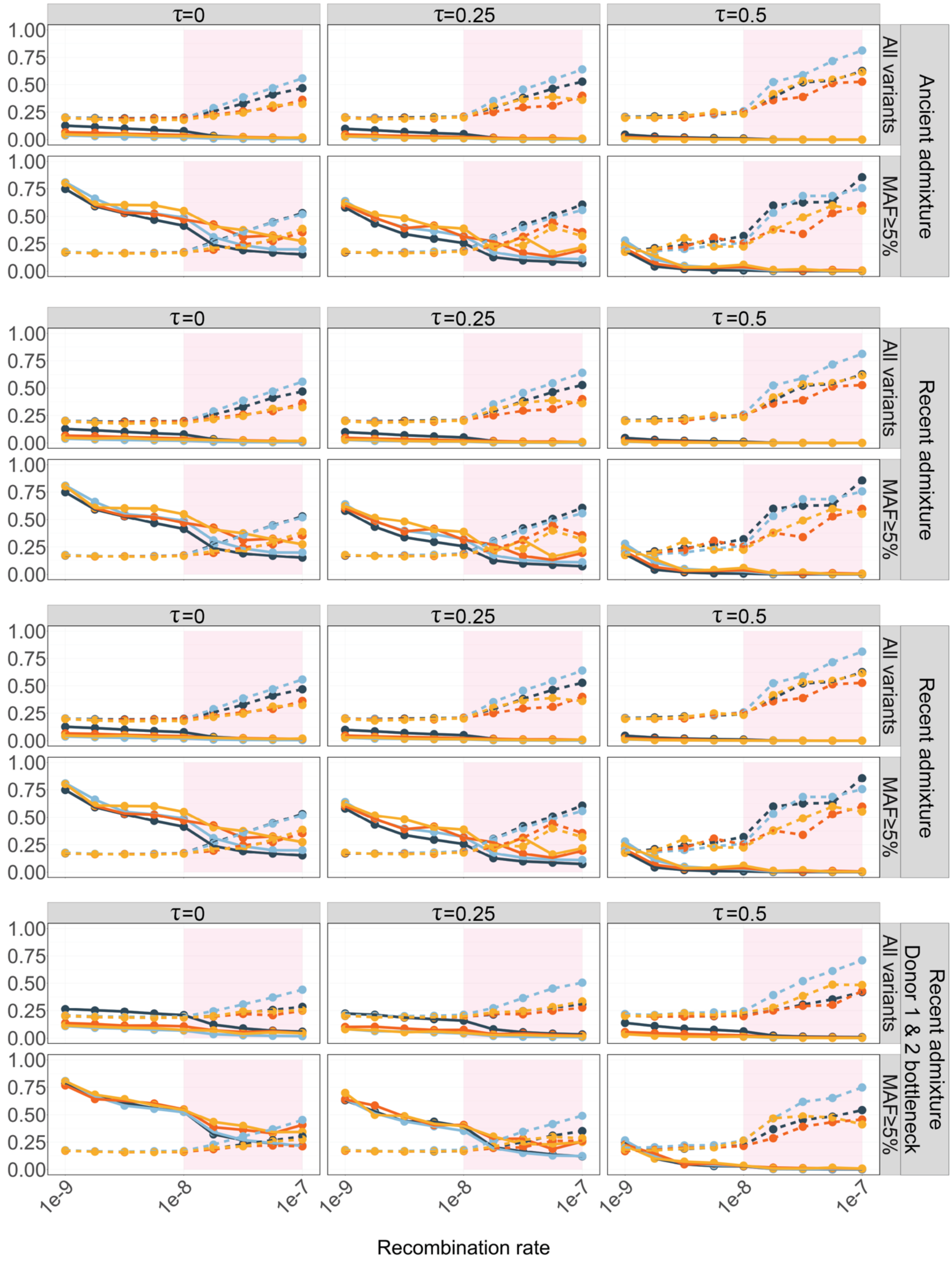

### Figure S5)

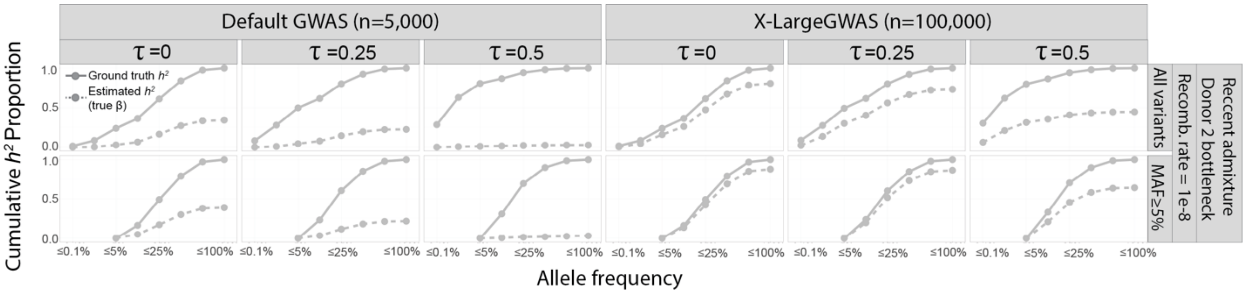
Default GWAS (n=5,000)
X-LargeGWAS (n=100,000)
τ =0
τ =0.25
τ =0.5
Ground truth h²
Estimated h² (true β)
Cumulative h² Proportion
Allele frequency
≤0.1%
≤5%
≤25%
≤100%
All variants
MAF≥5%
Recomb. rate = 1e-8
Donor 2 bottleneck
Reccent admixture

**Supplementary Tables**

**Table S1. List of UKBiobank phenotypes with GWAS summary statistics under each UKB-PPP panel.**

| | UKB-PPP panel | | | |
|---|---|---|---|---|
| | Inflammation | Cardiometabolic | Neurology | Oncology |
| UKBiobank GWAS phenotypes | CRP (C-reactive protein)<br>WBC (white blood cell count)<br>Neutrophil<br>Lymphocyte | LDL<br>HDL<br>Cholesterol<br>Triglycerides<br>Glucose<br>BMI<br>Coronary heart disease<br>Hypertension<br>Stroke<br>T2D<br>Obesity | Alzheimer<br>Parkinson's<br>Multiple sclerosis<br>Epilepsy<br>Depression<br>Schizophrenia | Breast cancer<br>Prostate cancer<br>Colorectal cancer<br>Lung cancer<br>Ovarian cancer<br>Skin cancer<br>Larynx throat cancer<br>Tongue cancer<br>Stomach cancer<br>Kidney cancer<br>Colon cancer |

**Table S2. Number of variants from each UKB-PPP panels and number of variants from UKBiobank GWAS summary statistics.**

| | UKB-PPP panel | | | | | | | |
|---|---|---|---|---|---|---|---|---|
| | Inflammation | | Cardiometabolic | | Neurology | | Oncology | |
| | Panel I | Panel II | Panel I | Panel II | Panel I | Panel II | Panel I | Panel II |
| # of analytes | 368 | 369 | 369 | 367 | 367 | 367 | 368 | 368 |
| # of pQTL significant variants | 619,076 | | 594,359 | | 476,932 | | 425,081 | |
| # of GWAS significant variants | 138,118 | | 177,370 | | 22,992 | | 33,318 | |
| # of overlap between pQTL and GWAS significant variants | 69,047 | | 60,923 | | 12,153 | | 504 | |
